# Inhibition of the serine/threonine kinase BUB1 reverses taxane resistance in prostate cancer

**DOI:** 10.1101/2023.05.05.539598

**Authors:** Martinez Maria Julia, Lyles Rolando DZ, Peinetti Nahuel, Grunfeld Alex Michael, Burnstein Kerry L

## Abstract

**Background:** Men with incurable castration resistant prostate cancer (CRPC) are typically treated with taxanes; however, drug resistance rapidly develops. Thus, overcoming taxane resistant PC is a major clinical need. We previously identified a seven gene network in aggressive CRPC, which includes the mitotic serine threonine kinase *BUB1*, a major regulator of the spindle assembly checkpoint (SAC). Alterations in mitotic kinases (and SAC malfunction) are associated with advanced PC and taxane resistance development and thereby represent potential vulnerabilities.

**Methods:** We evaluated BUB1 expression in publicly available data sets and in existing and newly generated taxane resistant PC cells. The effects of BUB1 depletion on the growth of a panel of PC and non-tumorigenic prostate epithelial cells was determined. We examined the capacity of pharmacologic inhibition of BUB1 kinase to reverse taxane-resistant PC growth. We evaluated the role of the prevalent androgen receptor variant AR-V7, in regulating BUB1 expression and taxane resistance.

**Results:** BUB1 mRNA was over-expressed in PC, metastatic castration resistant prostate cancer (mCRPC) and in tumors of patients treated with taxane-based chemotherapeutics compared to benign prostate tissue. Furthermore, BUB1 levels were elevated in taxane resistant PC cell lines compared to their sensitive counterparts. BUB1 depletion decreased growth of CRPC cells through delayed mitosis but did not affect proliferation of androgen dependent (ADPC) or non-tumorigenic prostate epithelial cells. Furthermore, BUB1 inhibition with the specific kinase inhibitor, BAY1816032, re-sensitized taxane resistant CRPC cells to the clinically used drugs, docetaxel and cabazitaxel. Consistent with AR-V7 regulation of BUB1, we also found that AR-V7 was elevated in taxane resistant CRPC cells. Moreover, ectopic expression of AR-V7 in CRPC cells that lack this protein resulted in increased BUB1 and conferred docetaxel resistance. BUB1 pharmacologic inhibition in combination with taxanes sensitized AR-V7 expressing CRPC cells to docetaxel treatment.

**Conclusion:** These data support BUB1 as an exploitable and therapeutically tractable vulnerability in taxane resistant CRPC including in AR variant driven CRPC.

## Introduction

Androgen deprivation therapy (ADT) is the standard of care for high-risk localized or advanced / metastatic prostate cancer (PC)(1, 2). Although ADT potently decreases tumor growth, within 24-36 months the disease typically progresses to incurable castration resistant prostate cancer (CRPC)(1, 3). After ADT fails, patients with CRPC or metastatic CRPC (mCRPC) are commonly treated with docetaxel. The semisynthetic taxane, cabazitaxel, is a second generation taxane, which is available for those mCRPC patients whose disease progresses after docetaxel treatment. However, within months, tumors of patients treated with cabazitaxel become refractory to taxanes leaving no other life-prolonging options for these patients(1, 4). Both docetaxel and cabazitaxel block microtubule dynamics, leading to prolonged mitotic arrest through sustained activation of the spindle assembly checkpoint (SAC) and eventually triggering cell death(5). Taxane-resistance can develop through a variety of mechanisms, including altered structure of microtubules, aberrant G2/M transition, defects in the SAC or premature mitotic exit(6, 7). While there is currently no consensus as to whether androgen receptors (AR) and/or AR splice variants (AR-Vs) participate in taxane responsiveness and resistance development (8–10). AR-V7 expression in circulating tumor cells is associated with poorer outcome in taxane-treated mCRPC patients (11–13).

Using an unbiased systems biology approach, we previously identified an AR-V7 driven clinically relevant gene network that is composed of cell cycle and mitotic genes(14). The gene network members are overexpressed in PC, and elevated expression of these genes strongly predicts decreased patient survival in PC patient cohorts. The gene set includes the serine threonine kinase BUB1, an essential component of the SAC(14). After activation of the SAC, BUB1 is recruited to unattached kinetochores and functions as a scaffold protein required for kinetochore localization of other checkpoint proteins. As a kinase, BUB1 phosphorylates Histone H2A (Thr120) at mitotic centromeres to recruit Shugoshin 1 and 2, which protects centromeric cohesion and recruits Aurora B to the chromosomal passenger complex (CPC) to correct errors in kinetochore-microtubules (KT-MT) attachment(15, 16). Higher BUB1 mRNA expression is associated with poorer clinical outcome in several types of cancer including liver(17), bladder(18), and neuroblastoma(19); however, BUB1’s possible roles in PC and chemotherapeutic resistance remains understudied.

Since SAC alterations are one of the mechanisms underlying the development of taxane resistance and because BUB1 is up-regulated in mCRPC, we evaluated the potential of BUB1 to serve as a therapeutic target in clinically relevant advanced PC, including CRPC and taxane resistant (docetaxel and cabazitaxel) cell lines. Inhibition of BUB1 with the specific kinase inhibitor, BAY-1816032, sensitized resistant cells to taxane treatment (docetaxel or cabazitaxel), through mitosis delays and increased apoptosis in resistant cell lines. Moreover, ectopic AR-V7 expression increased levels of BUB1 protein and promoted resistance to docetaxel. Our study supports BUB1 as a potential therapeutic target for advanced PC and supports BUB1 inhibition as a tractable mechanism to overcome taxane-resistance in mCRPC.

## Material and Methods

### Analysis of human samples datasets

The TCGA-PRAD was downloaded from to the Xena Functional Genomics Explorer (http://xena.ucsc.edu/). MSKCC (Cancer cell, 2010), SU2C (Cell, 2019) and Hutchinson (Nat med, 2016) data sets were accessed and downloaded from cBioportal (cbioportal.org). The GSE35988 (Grasso et al, 2012) and GSE32269 (Cai et al, 2012) data sets were downloaded from from GEO portal (https://www.ncbi.nlm.nih.gov/gds). The BUB1 expression values for all datasets were presented as log2-transformed signal intensity ratios. To build Kaplan-Meier curves, patient samples were stratified in quartiles based on BUB1 expression and disease-free curves were built comparing patients in the top quartile (high BUB1) compared to patients in the bottom quartile (low BUB1).

### Cell culture and reagents

Human PC cell lines RWPE-1, LNCaP, C4-2B, 22Rv1 (CRL-2505), PC3 and DU145 were obtained from American Type Culture Collection (ATCC, Manassas, VA, USA). BPH-1, PC3 and DU145 taxane sensitive (TxS), taxane resistant (TxR) and cabazitaxel (cabR) resistant cells were gifts from fellow scientists (see acknowledgments). All cell lines were negative for mycoplasma, bacteria, and fungi contamination. BPH-1, LNCaP, 22Rv1, DU145 and PC3 cells were cultured in RPMI (Corning, New York, USA) supplemented with penicillin (100 IU/mL), streptomycin (100 μg/mL), 2 mM L-glutamine (Life Technologies, Carlsbad, CA, USA), and 10% fetal bovine serum (FBS) (Hyclone, San Angelo, TX, USA). C4-2B cells were cultured in DMEM (Corning), supplemented as detailed above. RWPE-1 cells were cultured in Keratinocyte SFM (Thermo Fisher Scientific, Waltham, MA, USA) supplemented with bovine pituitary extract and EGF (Thermo). 22Rv1, PC3 and DU145 TxR and cabR cells were cultured with 10 nM docetaxel (Sigma-Aldrich, St. Louis, MO, USA) or cabazitaxel (Sigma Aldrich) respectively. For drug treatment assays, cells were cultured in 2 or 10% CSS. Cell cultures were maintained at 37°C in a humidified atmosphere of 5% CO2. BAY-1816032 (CT-BAY-1816032), was purchased from Chemietek (Indianapolis, IN, USA).

### Development of taxane resistant 22Rv1 cell lines

22Rv1 docetaxel (TxR) and cabazitaxel (cabR) resistant models were generated as previously published(20, 21). The development of each resistant cell line took approximately 6 months.

### Plasmids

The pLKO.1-shBUB1 (TRCN0000288618, Sigma Aldrich) plasmid against the coding region was used for knockdown experiments. The pLKO.1-shGFP, pLVX-EV and pLVX-AR-V7 plasmids were provided by fellow scientists (see acknowledgments). Cells transduced with pLVX or pLKO.1 plasmids were selected in 2.5 µg/ml of puromycin for 2 to 3 days.

### Evaluation of cell proliferation, mitosis and apoptosis by live cell imaging

For proliferation assays, PC3, DU145, TxS, TxR, cabR (5 000 cells per well), 22Rv1 TxS, TxR and cabR (10 000 cells per well) cells were seeded in 96-well plates (Corning) and incubated 37°C overnight. Cells were treated with different doses of BAY-1816032, docetaxel, cabazitaxel or their combination (in 10% CSS) and proliferation was measured by the Incucyte Zoom System (Essen Bioscience Inc., Ann Arbor, MI, USA). For apoptosis assays, at the time of treatment, cells were transfected with 1% (v/v) IncuCyte Caspase-3/-7 Apoptosis Assay Reagent (Essen Bioscience Inc.). Phase and green-fluorescent images were obtained at 10X every 4 h for 72 h and images were analyzed using the Incucyte Zoom software.

To assess mitosis duration, cells were seeded in 24-well plates at 20 000 cells per well and incubated at 37°C and imaged using the Incucyte Zoom System (Essen BioScience Inc). Cells were imaged every 20 minutes for 48 h after the start of incubation at 20X magnification. Individual cells (50 cells per group) were tracked from the beginning to end of mitosis.

### Evaluation of cell proliferation by Trypan blue exclusion assay

For BUB1 knockdown experiments cells were seeded in 24 well plates at 10 000 cells per well (RWPE-1, BPH-1), 20 000 cells per well (LNCaP, 22Rv1, PC3 and DU145) or 40 000 cells per well (C4-2B). For drug treatment assays, PC3 and DU145 TxS, TxR and cabR (20 000 cells per well) and 22Rv1 TxS, TxR and cabR (40 000 cells per well) were seeded in 24-well plates. Cells expressing pLVX-EV and pLVX-AR-V7 (30 000 cells per well) were seeded in 24-well plates. 24 h after seeding, cells were washed with PBS and treated with different doses of BAY-1816032, docetaxel, cabazitaxel or their combination for 72 h. At the indicated time points cells were trypsinized (Corning) and mixed with trypan blue (Gibco, Billings, MT, USA) in 1 to 1 ratio, and live cells were counted using the Countess II (Thermo Fisher).

### Soft Agar Assays

Soft agar assays were performed as previously described(22). DU145 and PC3 TxS, TxR and cabR cells (200 000 per 35 mm plate) were treated (vehicle, BAY-1816032 0.5 µM, docetaxel 10 nM, cabazitaxel 10 nM or their combination) at the time of seeding and incubated for 3 weeks. Media was changed once every week. Images were taken with microscope and colonies were counted using Image J software.

### RNA isolation and RT-qPCR

Total RNA was harvested using TRIzol (Thermo Fisher) and purified using Direct-zol RNA MiniPrep Plus reagent (Zymo Research, Irvine, CA, USA), according to the manufacturer’s protocol. Total RNA (0.5-1 μg) was reverse-transcribed using a High-Capacity cDNA Reverse Transcription kit (Thermo Fisher Scientific). Real-time PCR was performed using 100 ng of cDNA and ABI StepOnePlus (Thermo Fisher). TaqMan probes from Thermo Fisher Scientific for *BUB1* (Hs01557695_m1) and *GAPDH* (HS99999905) were used. The comparative threshold cycle method was used to determine the relative expression of mRNA.

### Immunoblotting

Protein samples were harvested, and Western blots were performed as previously described(22). Cleaved PARP (9541), anti-rabbit (7074s) and anti-mouse (7076s) antibodies were purchased from Cell Signaling Technology (Danvers, MA, USA). Actin (sc-1616), Cyclin A (sc-751) and anti-goat (sc-2354) were purchased from Santa Cruz Biotechnology (Santa Cruz, CA, USA). BUB1 (ab195268) antibody was purchased from Abcam (Cambridge, UK). GAPDH (MA5-15738) was purchased from Invitrogen (Waltham, MA, USA).

### Statistical analysis

Data were tested for normality (Shapiro–Wilk test) and homogeneity of variances (Levene’s test). When assumptions were met, data were tested for significance (p < 0.05) using a two-tailed Student’s t-test (two groups) or analysis of variance (ANOVA; three or more groups). Otherwise, non-parametric statistical analyses were used: Mann–Whitney’s test (two groups) and Kruskal– Wallis (unpaired, three or more groups) or Friedman (paired, three or more groups). Log-rank test was used to test the significance of differences in Disease Free curves. Results are expressed as means ± SEM, and p < 0.05 was considered significant. Combinatory index (CI) was calculated using the CompuSyn software (ComboSyn, Inc, Paramus, NJ, USA). Statistical analyses and graph plotting were carried out using GraphPad Prism software version 8.4 (San Diego, CA, USA).

## Results

### BUB1 is up-regulated in localized prostate cancer and metastatic castration resistant prostate cancer

Analysis of the Grasso(23) and TCGA-PRAD PC patient data sets revealed that BUB1 expression was significantly higher in primary PC compared to benign tissue. Furthermore, the Grasso, MSKCC (2010), Hutchinson (2016) and Cai(24) data sets demonstrated that BUB1 expression was significantly upregulated in mCRPC compared to localized PC (Figure 1 a). Survival metrics in three independent patient sample cohorts, showed that patients harboring high-BUB1 expressing tumors had decreased overall- and disease-free survival compared to patients with low-BUB1 expression (Figure 1 b). Thus, BUB1 up-regulation is associated with PC progression and a more rapid decline in patient survival.

**Figure 1.**
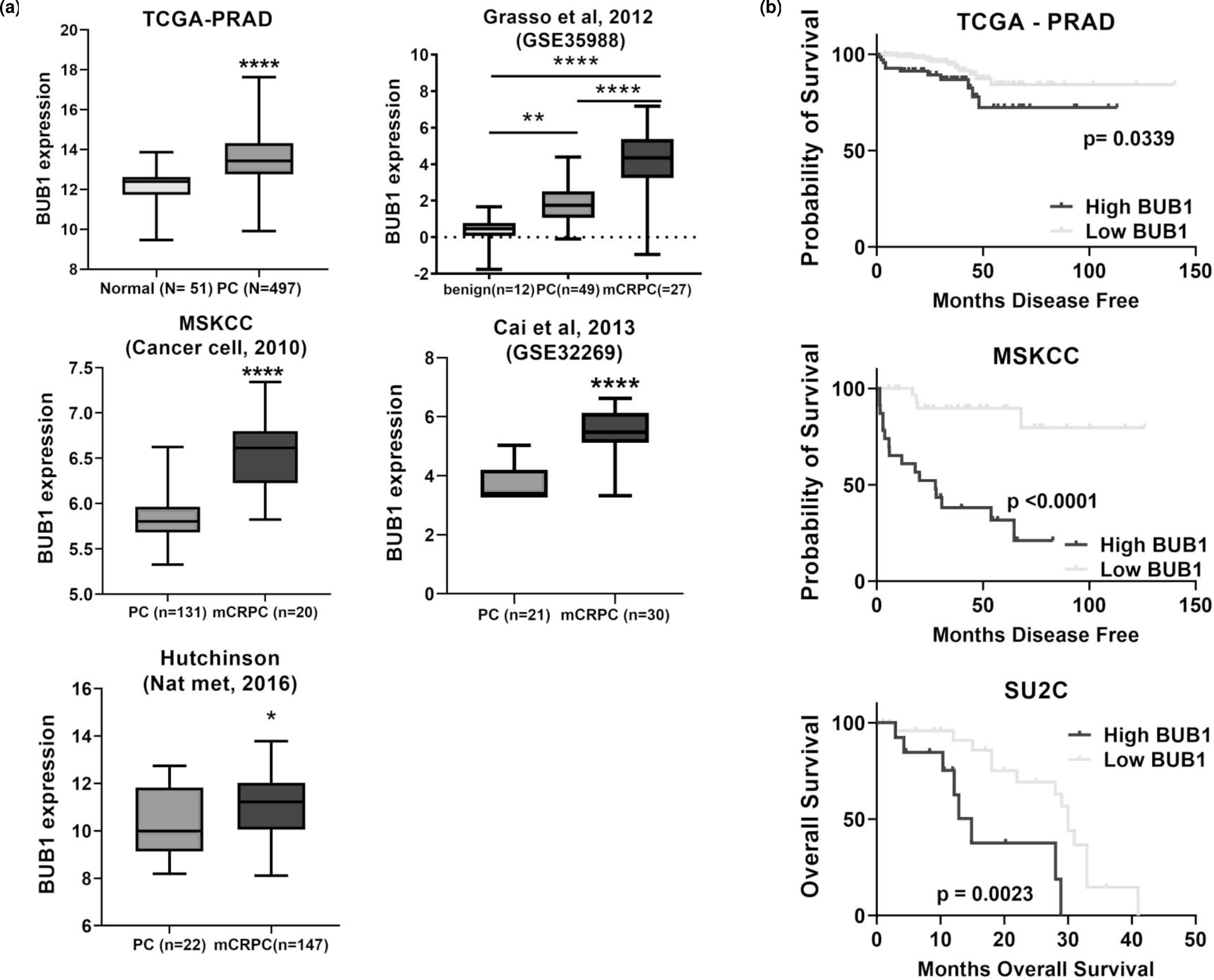
BUB1 is up-regulated in PC and CRPC human tumor samples and is prognostic for poor outcome. **(a)** TCGA-PRAD was downloaded from UCSC Xena (http://xena.ucsc.edu/) and MSKCC and Hutchinson were downloaded from cBioportal and Grasso et al, 2012 and Cai et al, 2013 were downloaded from GEO portal (https://www.ncbi.nlm.nih.gov/gds). (Student’s t test, ** p < 0.01, *** p < 0.001, **** p < 0.0001). **(b)** Kaplan-Meir curves for overall survival (SU2C) and disease-free survival (MSKCC, Cancer Cell, 2010) in patients with high BUB1 (top quartile) expression compared to patients with low BUB1 (bottom quartile) expression. Graphs were elaborated using GraphPad (LogRank test, * p < 0.05, **** p < 0.0001).

### BUB1 depletion decreased growth of CRPC but not androgen dependent PC or non-tumorigenic prostate epithelial cells

While BUB1 depletion did not significantly affect growth of non-tumorigenic RWPE-1 or BPH-1 cell lines or androgen dependent LNCaP cells, BUB1 knockdown significantly decreased proliferation of CRPC cell lines (Figure 2 a). BUB1 was expressed (mRNA and protein) in all evaluated cell lines (Supplementary Figure 1), thus the lack of growth inhibition after BUB1 depletion in BPH1, RWPE-1 and LNCaP cells suggests that BUB1 selectively promotes CRPC cell growth. Moreover, the differential effect of BUB1 depletion on growth of CRPC cells vs androgen dependent or benign prostate cells was not due to differences in knockdown efficiency (Supplementary Figure 2 a) and was associated with significant delay in mitosis progression in CRPC cells and decreased expression of Cyclin A levels in CRPC cells (Figure 2 b, Supplementary Figure 2 b). However, no significant increase in the percent of dead cells or activation of caspase dependent apoptosis was observed after BUB1 depletion in CRPC cells (Supplementary Figure 2 c and d). These results suggest that BUB1 specifically enhanced CRPC proliferation through mitosis promotion.

**Figure 2.**
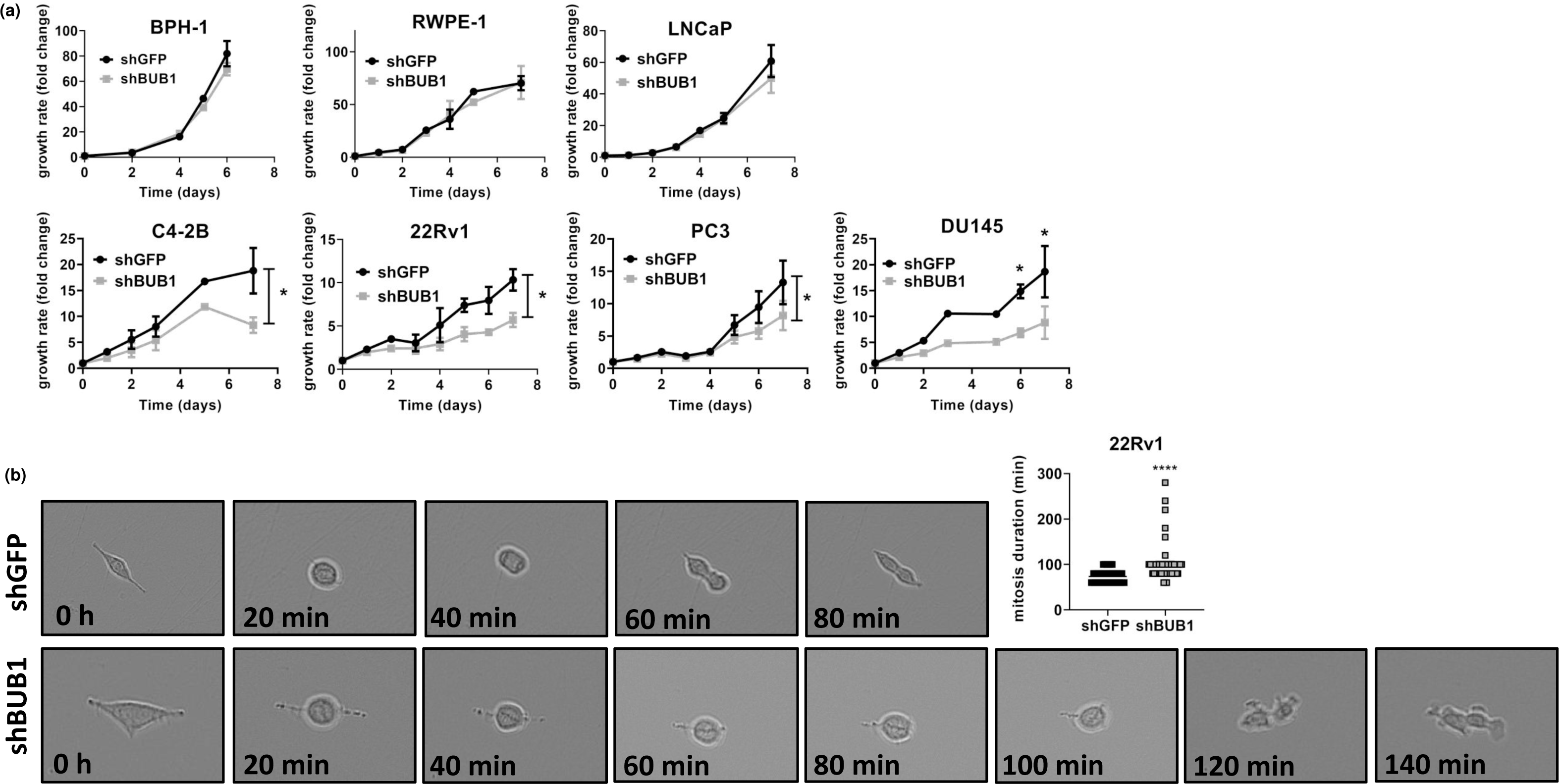
BUB1 knockdown decreased CRPC cell proliferation through cell cycle inhibition. **(a)** Proliferation of the indicated cell lines was measured by trypan blue exclusion after BUB1 depletion at different time points. Graphs represent the average of two (DU145 and BPH-1), three (RWPE-1), four (C4-2B and PC3) or five (22Rv1 and LNCaP) independent experiments performed in quadruplicate ± SEM. The area under the curve was calculated using GraphPad Prism. For DU145 cells, significant differences between shGFP and shBUB1 were calculated for each time point (Mann-Whitney test, * p < 0.05). **(b)** Mitosis duration was evaluated in 22Rv1 cells after BUB1 depletion using an Incucyte Zoom. Images were taken every 20 min and mitosis duration was quantified. Graph represents the quantification (50 cells per group) of a representative experiment from two independent experiments. (Unpaired Student T test, **** p < 0.0001).

### BUB1 expression is upregulated in taxane resistant CRPC cells

Given BUB1’s central role in SAC and mitosis progression and the fact that mitotic errors can promote taxane resistance in a variety of cancers (6, 7, 25, 26), we investigated BUB1 mRNA expression in patients with mCRPC who were treated or not with taxane-chemotherapy. BUB1 mRNA was significantly up-regulated in mCRPC samples from patients treated (67 patients) compared to non-taxane-treated patients (45 patients) in the Hutchinson dataset (Figure 3 a). Similarly, in the SU2C data set, BUB1 mRNA expression was significantly higher in metastatic samples from patients who received taxanes (77 patients) compared to non-treated patients (122 patients) (Figure 3 a). In contrast, the expression of BUB1B, a closely related serine/threonine kinase that is also part of the SAC, was not significantly different in taxane-treated compared to naïve patients (Supplementary Figure 3 a).

**Figure 3.**
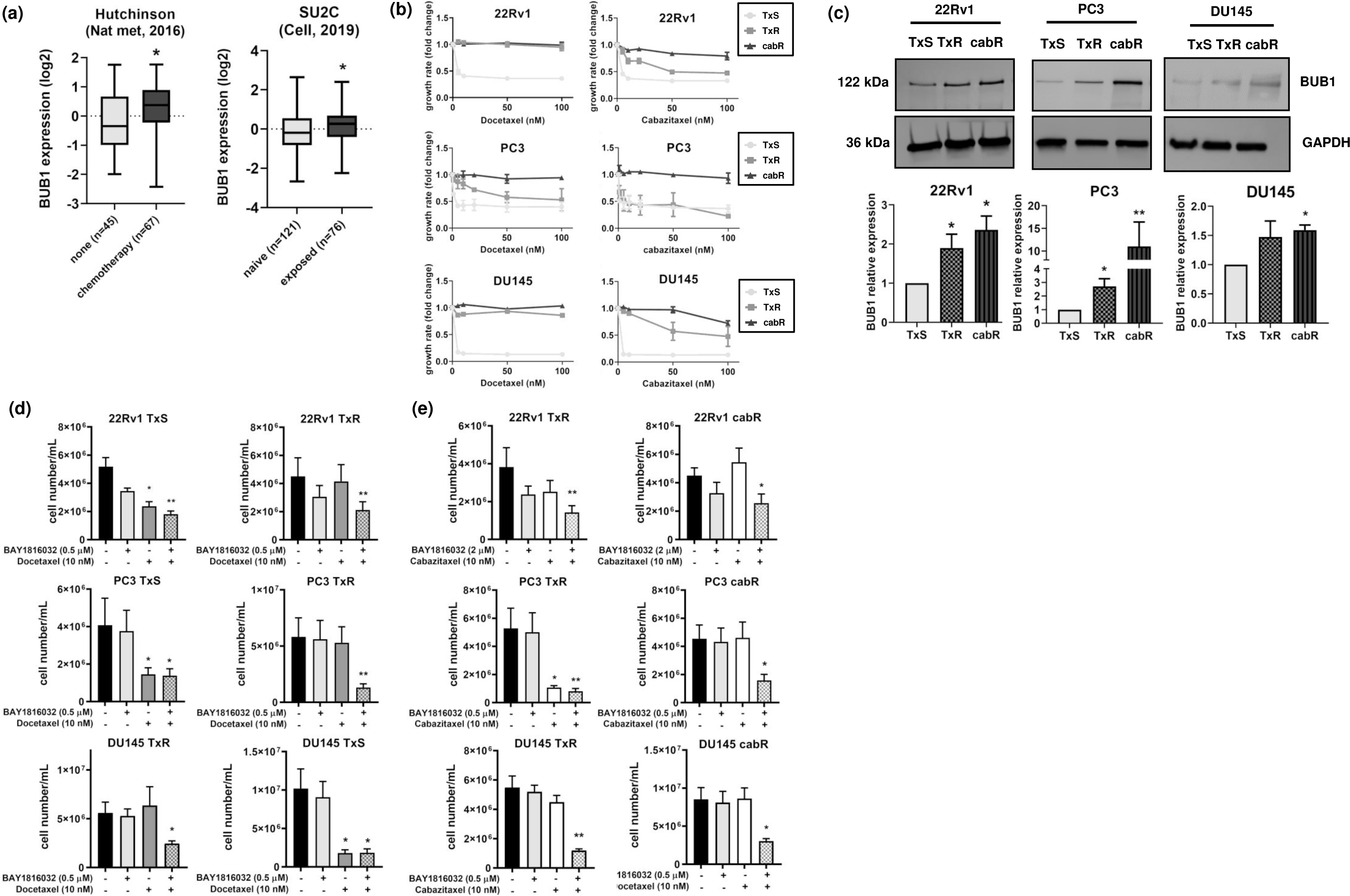
BUB1 inhibition re-sensitized taxane resistant PC cells to docetaxel or cabazitaxel. **(a)** BUB1 expression in metastatic samples from patients treated with chemotherapy (n=67) or non-treated (n=45) (Hutchinson (Nat Med, 2016)), or with taxanes (n=77) was compared to untreated (naïve) patients (n= 121) (SU2C, Cell 2015). (Data were downloaded from cBioportal; Student’s t test, * p < 0.05). **(b)** The indicated docetaxel sensitive (TxS), resistant (TxR) and cabazitaxel resistant (cabR) cell lines were treated with 0 – 100 nM doses of docetaxel or cabazitaxel for 72 h. Graphs represent the average of two to four independent experiments performed in quadruplicate ± SEM. **(c)** BUB1 protein levels in the indicated TxS, TxR and cabR cell lines were evaluated by Western Blot. Graphs represent the quantification of five independent experiments ± SEM. (Kruskal-Wallis test, * < p 0.05, ** p < 0.01). 22Rv1, PC3 and DU145 **(d)** TxS and TxR or **(e)** TxR and cabR cells were treated with BAY-1816032 (0.5 µM or 2 µM), docetaxel (10 nM) or cabazitaxel (10 nM) individually or in combination for 72 h. Cell proliferation was measured by trypan blue exclusion assay. Graphs represent the average of five independent experiments performed by quadruplicate ± SEM. (Kruskal-Wallis test, * < p 0.05, ** p < 0.01).

To explore the role of BUB1 in taxane resistance, we developed resistant CRPC cell lines by culturing 22Rv1 with increasing concentrations of docetaxel or cabazitaxel using a previously described method(20, 21). The resulting 22Rv1 TxR and cabR cells were validated by challenging the resistant cell lines with docetaxel and cabazitaxel and comparing cell proliferation to previously established DU145 and PC3 TxR and cabR models (20, 21). While none of the TxR models were affected by docetaxel as anticipated, these cell lines were inhibited by cabazitaxel (i.e., cross resistance was not observed). None of the cabR cell lines were inhibited by 500 nM docetaxel (data not shown) or cabazitaxel (Figure 3 b). Supplementary table 1 shows the IC_50_ values for the TxS, TxR and cabR cell lines treated with docetaxel or cabazitaxel respectively. We found that BUB1 protein levels were significantly higher in TxR and cabR cell lines compared to their sensitive counterparts (Figure 3 c). BUB1 mRNA expression followed a similar trend, showing higher levels in the resistant models (Supplementary Figure 3 b).

### BUB1 inhibition rescues taxane sensitivity through mitosis delay and apoptosis

To explore the effect of BUB1 kinase inhibition on the proliferation of taxane resistant CRPC cells, we combined BAY-1816032 with docetaxel or cabazitaxel in TxS, TxR and cabR cell lines. First, we performed BAY-1816032 dose response experiments (Supplementary Figure 3 c) to select sub-optimal drug concentrations (does not significantly inhibit cell proliferation) that were then used for combination drug treatment experiments. We selected BAY-1816032 concentrations below 1 µM for PC3 and DU145 and up to 2 µM for the 22Rv1 TxS, TxR and cabR cells.

Notably, BAY-1816032 plus taxane synergistically [combination index (CI) < 1] inhibited TxR and cabR cell proliferation (Figure 3 d and e, Supplementary Figure 4 a and b). Combination treatments also significantly reduced TxR and cabR anchorage independent growth (Supplementary Figure 4 c and d).

BUB1 regulation of SAC and CPC in mitosis avoids chromosome segregation errors, that could lead to chromosomal instability and aneuploidy. In line with this function, the combination of BAY-1816032 and either docetaxel or cabazitaxel significantly prolonged mitosis duration in PC3 taxane resistant cells (Figure 4 a – d). Since, prolonged mitotic arrest could lead to cell death, we evaluated apoptosis activation after dual treatment. BAY-1816032 in combination with docetaxel or cabazitaxel significantly increased apoptosis, as evidenced by increased caspase -3/-7 activity TxR and cabR cells (Figure 4 e and f and Supplementary Figure 4 e and f). Moreover, BUB1 inhibition in combination with docetaxel or cabazitaxel decreased expression of the cell cycle marker, Cyclin A and increased levels of the pro-apoptotic marker cl-PARP in 22Rv1 TxR or cabR cells (Figure 4 g and h). Thus, BUB1 kinase inhibition re-sensitized TxR and cabR PC cells to taxanes by prolonging mitosis duration and activating apoptosis.

**Figure 4.**
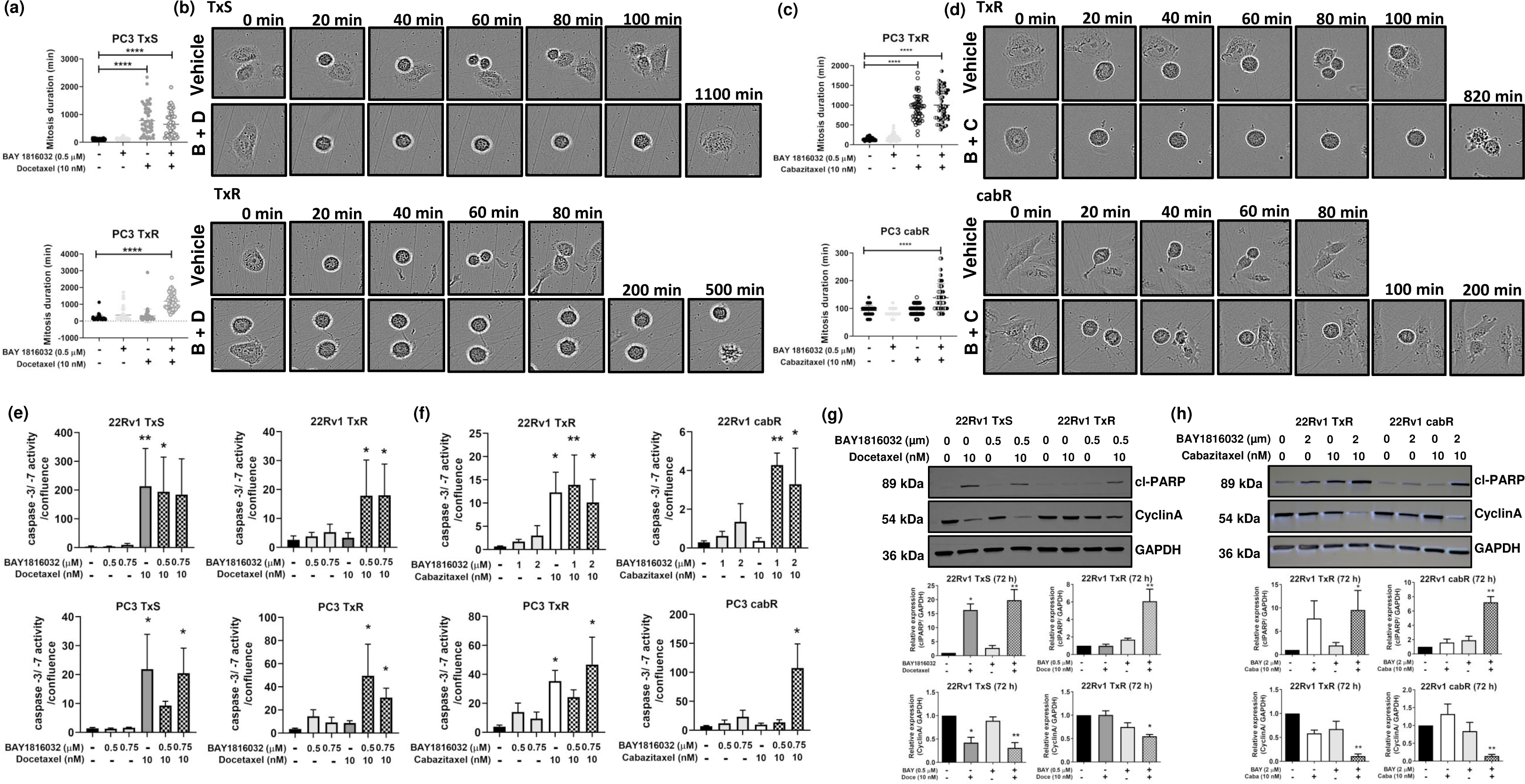
BUB1 inhibition sensitized TxR and cabR cells to docetaxel or cabazitaxel treatment through mitosis delay and apoptosis induction. Mitosis duration as described in Fig. 2 legend was evaluated in PC3 **(a)** TxS and TxR or **(c)** TxR and cabR cells after treatment with BAY-1816032 (0.5 µM), docetaxel (10 nM), cabazitaxel (10 nM) or their combination for 24 h. Graph represents the quantification (50 cells per group) of a representative experiment from two independent experiments (Unpaired Student T test, **** p < 0.0001). PC3 **(b)** TxS and TxR or **(d)** TxR and cabR, vehicle or BAY-1816032 plus taxanes, representative images are shown. 22Rv1 and PC3 **(e)** TxS and TxR or **(f)** TxR and cabR cells were treated with BAY-1816032 (0.5 or 0.75 µM), docetaxel (10 nM), cabazitaxel (10 nM) or their combination for 72 h. Cells were transfected with caspase 3/ 7 dye and caspase activity was measured by Incucyte Zoom. Graphs represent the average of four independent experiments performed in quadruplicate ± SEM. Kruskal-Wallis test, * p < 0.05, ** p < 0.001. cl-PARP and CyclinA levels were quantified in 22Rv1 **(g)** TxS and TxR or **(h)** TxR and cabR by Western Blot after treatment with BAY-1816032 (0.5 or 2 µM), docetaxel (10 nM), cabazitaxel (10 nM) or their combination for 72 h. Graphs represent the quantification of four independent experiments ± SEM. (Kruskal-Wallis test, * p < 0.05, ** p < 0.01). Blots are representative of four independent experiments.

### AR-V7 promoted increased BUB1 expression leading to docetaxel resistance

Since we previously identified BUB1 as part of an AR-V7 driven network(14) and AR-V7 is associated with the development of taxane resistance(8, 9, 27), we measured AR-V7 expression in 22Rv1 TxS and TxR cells. AR-V7 levels were higher in 22Rv1 TxR and cabR compared to TxS cells (Figure 5 a). Notably, no significant changes were observed in full length AR levels (Supplementary Figure 5 a). Moreover, we observed that BUB1 inhibition in combination with docetaxel or cabazitaxel significantly reduced AR and AR-V7 in 22Rv1 TxR and cabR cells (Supplementary Figure 5 b). Ectopic expression of AR-V7 in two AR-Vs null CRPC cells lines, C4-2B and PC3, resulted in increased BUB1 protein levels (Figure 5 b). Moreover, AR-V7 expression increased resistance to docetaxel treatment (Figure 5 c). BUB1 inhibition in combination with docetaxel significantly reduced PC3-AR-V7 cell proliferation (Figure 5 d). These results indicate that the AR-V7/BUB1 axis can participate in taxane resistance.

**Figure 5.**
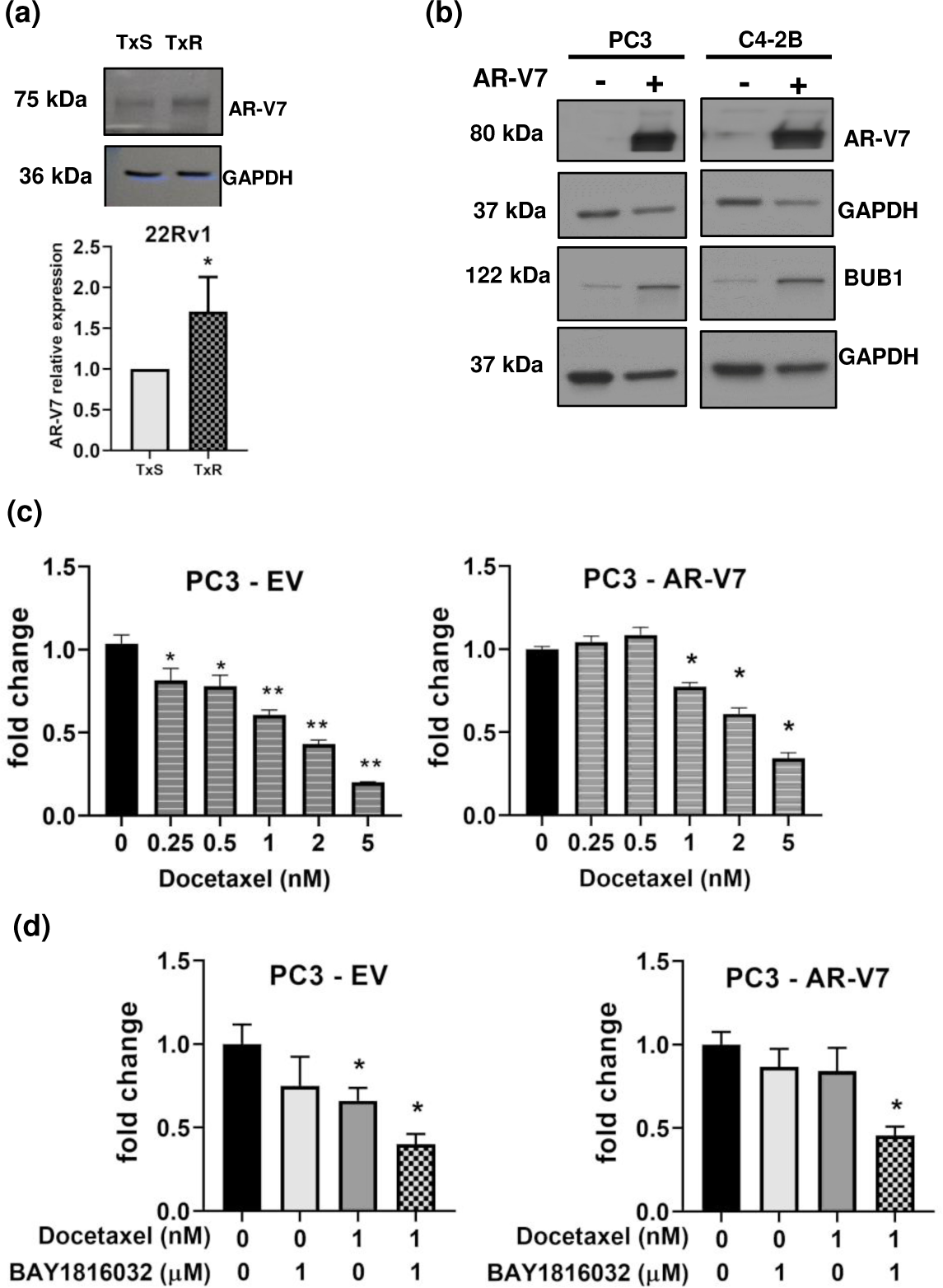
Ectopic AR-V7 expression increased BUB1 levels and resistance to taxanes in CRPC cells. **(a)** AR-V7 expression was measured in 22Rv1 TxS and TxR cells by Western Blots. Graph represents the quantification of four independent experiments ± SEM. (Mann-Whitney test, * p < 0.05). **(b)** PC3 and C4-2B cells were transfected with pLVX-Ev or pLVX-AR-V7 and expression of BUB1 and AR-V7 was evaluated by Western Blot. Blots are representative from two independent experiments. **(c)** PC3 EV or AR-V7 cells were treated with different doses of docetaxel for 72 h and cell viability was measured by Trypan blue exclusion assay. Graphs represent the average of four independent experiments performed in triplicates ± SEM. (Kruskal-Wallis test, * p < 0.05, ** p < 0.01, *** p < 0.001). **(d)** PC3 EV or AR-V7 cells were treated with BAY-1816032 (1 µM), docetaxel (1 nM) or their combination for 72 h and cell viability was measured by Trypan blue exclusion assay. Graph represents two independent experiments performed in triplicates ± SEM. (Kruskal-Wallis test, * p < 0.05)

## Discussion

This study supports BUB1 as a therapeutic target in taxane resistant CRPC. We showed that BUB1 mRNA is elevated in PC and mCRPC patients and that high BUB1 expression is evident in mCRPC patients after taxane treatment. BUB1 depletion specifically decreased CRPC proliferation, without significantly affecting ADPC or non-tumorigenic prostate epithelial cell growth. BUB1 kinase inhibition was sufficient to restore taxane sensitivity. Moreover, ectopic expression of AR-V7 increased BUB1 levels and was associated with reduced sensitivity to docetaxel. BAY-1816032 in combination with docetaxel, significantly decreased proliferation of AR-V7 overexpressing cells.

BUB1 specifically induced CRPC proliferation by promoting mitosis progression. In agreement with our findings, BUB1 promotes cell proliferation of several different cancer cell lines such as osteosarcoma, ovarian and hepatocellular (17, 28, 29). Moreover, BUB1 pro-proliferative effect has been associated with TGF-β pathway activation (17, 29). In a lung cancer cell line, BUB1 kinase activates TGF-β pathway through SMAD-2 and -3 and PI3K/AKT phosphorylation (30–32).

Identification of synergistic drug combinations is a strategy to overcome chemotherapy resistant CRPC. Since the mechanism of action of taxanes include the disruption of spindle dynamics, it has been hypothesized that targeting mitosis surveillance mechanisms, such as SAC or the CPC, could increase the efficacy of taxanes (25, 33, 34). This hypothesis is supported by recent findings demonstrating that high aneuploidy and CIN represent vulnerabilities in cancer and PC(26, 35–37), and that high CIN correlates with increased expression of mitotic kinases (including BUB1) and development of taxane resistant PC(26). Alteration of mitotic kinases such as PLK1, Msp1 and BUB1 have been observed in different taxane resistant cellular models including ovarian, breast and prostate cancer(25, 38–40). Furthermore, individual inhibition of different mitotic kinases such as AURKB, KIF20A and TOP2A results in significant reduction of cabazitaxel resistant cell proliferation *in vitro* and *in vivo*(41, 42). In agreement with these findings, we identified BUB1 mitotic serine threonine kinase but not the closely related kinase BUB1B, as being up-regulated in patients after receiving chemotherapy and in taxane resistant cell lines, positioning BUB1 as a potential vulnerability for advanced PC.

We showed that disruption of BUB1’s catalytic activity was sufficient to reverse the taxane resistant phenotype. Previous reports demonstrated that inhibition of BUB1 kinase activity only produces severe defects in mitosis progression after combination with taxanes in sensitive cell lines(43, 44),. Here we showed that BAY-1816032 plus taxanes resulted in synergistic cell growth inhibition not only in parental cells but importantly re-sensitized taxane resistant cells. Only minor effects on SAC have been reported after inhibition of BUB1 kinase activity(43, 44); suggesting that the re-sensitization observed after BAY-1816032 plus taxanes might be due to uncorrected MT-KT errors and not due to SAC disruption. Thus, targeting BUB1 kinase activity, by disrupting its role in CPC, might represent a vulnerability for taxane resistant CRPC cells. Our results support further study of BUB1 in taxane resistance development and the evaluation of the dual treatment in other clinically relevant models such as PDXs and *in vivo* taxane resistant models. Noteworthy, BAY-1816032 had to be administered orally twice a day in xenograft or orthotopic triple-negative breast cancer mouse models, suggesting poor bioavailability. Thus, to optimally test BUB1 inhibition in clinically relevant *in vivo* CRPC models, better inhibitors are needed.

While substantial evidence supports a role for AR-Vs in the development of chemotherapy (taxane) resistance in PC, there is still some controversy (6, 8, 10). Previous reports indicate that taxanes partially function by blocking AR but not AR-Vs nuclear translocation, and that increased AR-Vs expression (such as AR-V7 which undergoes microtubule-independent nuclear translocation) is a mechanism of taxane resistance(9, 45, 46). In addition, overexpression of AR-Vs, but not full-length AR, increases resistance to taxane treatment in sensitive PC cells(47). An association between AR-V7 expression and taxane response in patient samples has been shown in some but not all studies(48, 49). Tagawa and collaborators(11), demonstrated that patients enrolled in the TAXYNERGY clinical trial who harbored tumors without expression of AR-Vs (AR-V7 and AR-V567) responded better to taxane treatment. A recent meta-analysis including 36 studies, showed correlation between AR-V7 expression in CTCs and shorter overall survival in patients receiving taxane chemotherapy(13). Here we showed that 22Rv1 taxane resistant cell lines expressed higher AR-V7 levels than their sensitive counterparts. Moreover, ectopic AR-V7 expression in CRPC cells (without endogenous AR-V7 expression) increased BUB1 levels and reduced sensitivity to docetaxel treatment. Taxane sensitivity was restored after BAY-1816032 plus docetaxel in AR-V7 overexpressing cells. Thus, our results support a potential role of the AR-V7/BUB1 axis in taxane resistance development and suggest that BUB1 might represent a vulnerability in AR-V7-expressing taxane resistant CRPC tumors.

Taken together, our results showed synergy between BUB1 inhibition and taxane-based chemotherapy in docetaxel and cabazitaxel resistant cell models and support the study of BUB1 as a therapeutic target in taxane resistant mCRPC, for which no curative treatments currently exist.

## Authorship

MJM was responsible for experimental design, conducting the research, data analysis and writing. RDZL, NP and AG conducted research, analyzed data and assisted with writing. K.L.B. was responsible for experimental design, data analysis, providing funding, and writing.

## Acknowledgments

We gratefully acknowledge Dr. Evan Keller for providing PC3 and DU145 TxS, TxR and cabR cell lines. We thank Dr. Yan Dong for providing pLVX-EV and -AR-V7 constructs and Dr. Priyamvada Rai for providing pLKO.1-shGFP plasmid.

---

**Supplementary Figure 1.**
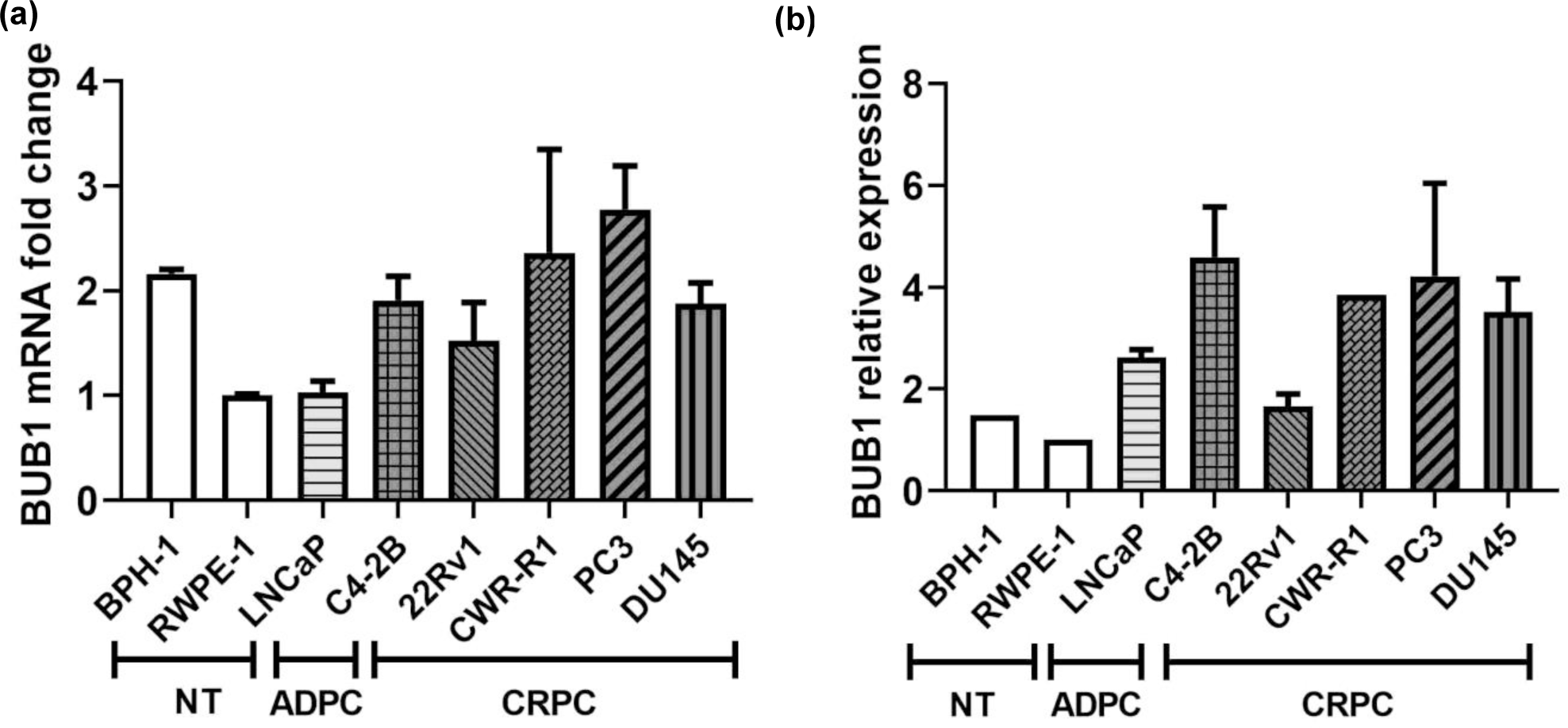
BUB1 expression in non-tumorigenic, PC and CRPC models. BUB1 **(a)** mRNA or **(b)** protein expression was evaluated in BPH-1, RWPE-1 (non-tumorigenic, NT), LNCaP (androgen dependent, ADPC), C4-2B, 22Rv1, CWR-R1, PC3 and DU145 (castration resistant, CRPC) by RTqPCR or Western blot. The graph represents the quantification of two (protein) or three independent (mRNA) experiments ± SEM.

**Supplementary Figure 2.**
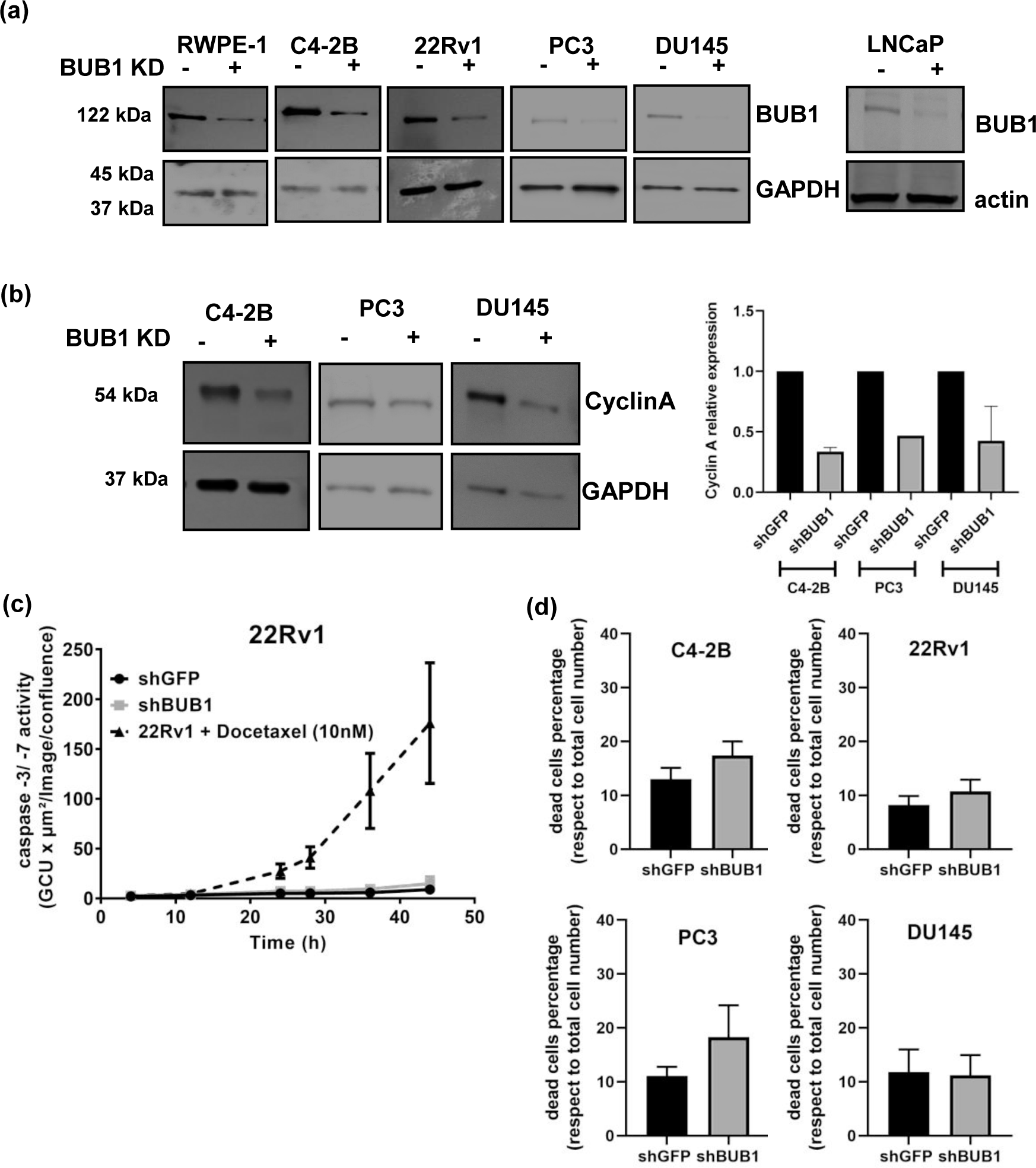
Non-tumorigenic or androgen-dependent PC cell growth was unaffected by Bub1b depletion. **(a)** BUB1 knockdown was evaluated by western blot in the indicated cell lines. **(b)** Expression of Cyclin A was measured 7 days after BUB1 knockdown by Western blot. Blots are representative of two independent experiments. **(c)** Caspase -3/ -7 activity was evaluated using the Incucyte Zoom in 22Rv1 cells after BUB1 depletion, 10 nM docetaxel was used as positive control. The graph represents one of two independent experiments performed in 8 technical replicates. **(d)** Dead cells were calculated as the percentage of trypan blue positive cells after 7 days for each treatment group. Graphs represent the average of two (DU145), four (C4-2B and PC3) or five (22Rv1) independent experiments performed in quadruplicate ± SEM.

**Supplementary Figure 3.**
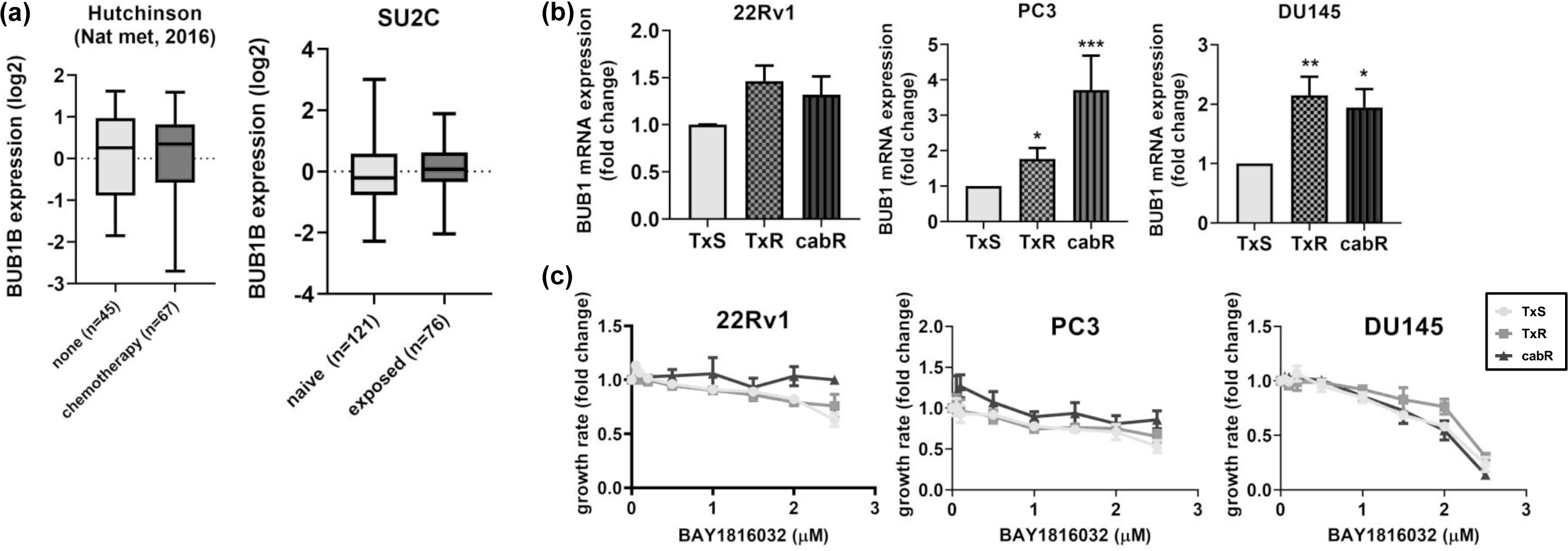
BUB1 expression was up-regulated in taxane resistant CRPC cell lines. **(a)** BUB1B expression in Hutchinson (Nat med, 2016) and SU2C (Cell, 2019) datasets (data was downloaded from cBioportal) was evaluated for metastatic samples that were treated with chemotherapy (n=67) or non-treated (n=45) (Hutchinson), exposed (n=77) to taxanes compared to naïve (n= 121) (SU2C, Cell 2015). (Student’s t test, * p < 0.05). **(b)** BUB1 mRNA expression was evaluated in taxane resistant PC cells by RTqPCR. Graphs represent the average of four to six independent experiments performed in triplicate ± SEM (Kruskal-Wallis test, * p < 0.05, ** p < 0.01). **(c)** Cells were treated with the indicated doses of BAY 1816032, for 72 h. Graphs represent the average of two to four independent experiments performed in quadruplicate ± SEM.

**Supplementary Figure 4.**
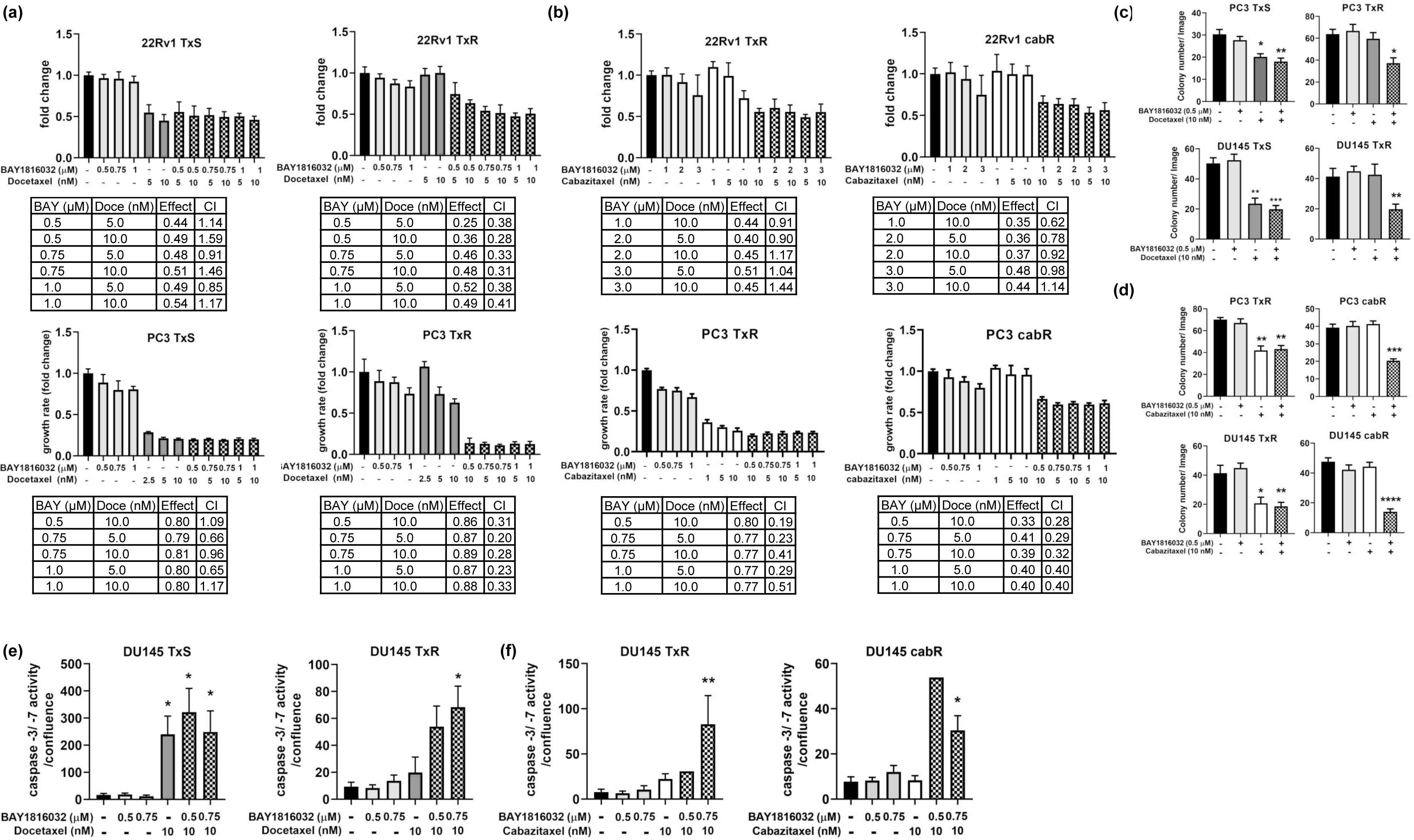
BUB1 inhibition in combination with docetaxel synergistically inhibited TxR cell proliferation. 22Rv1 and PC3. **(a)** TxS and TxR or **(b)** TxR and cabR cells were treated with BAY1816032 (0.5, 0.75, 1 and 2 µM), docetaxel (2.5, 5 and 10 nM), cabazitaxel (1, 5 and 10 nM) or their combination for 72 h. Proliferation was measured by Incucyte Zoom and combinatory index (CI) was calculated using Compusyn software. Graph represents a representative experiment from two independent experiments performed in octuplicate ± SEM. The table shows the CI for the different drug combinations. CI = 1 represents additivity, CI < 1 synergism, and CI > 1 antagonistic effect. DU145 **(c)** TxS and TxR or **(d)** TxR and cabR cells were treated with BAY1816032 (0.5 µM), docetaxel or cabazitaxel (10 nM) or the combination for 3 weeks and colony formation was measured by soft agar assay. Two images of each plate were taken, and colony number was quantified using Image J Software. Graphs represent the average of two independent experiments performed in triplicate ± SEM. DU145 **(e)** TxS and TxR or **(f)** TxR and cabR cells were treated with BAY1816032 (0.5 or 0.75 µM), docetaxel (10 nM), cabazitaxel (10nM) or their combination for 72 h. Cells were transfected with Incucyte caspase -3/ -7 dye and activity was measured using the Incucyte Zoom. Graphs represent the average of six independent experiments performed in quadruplicate ± SEM (Friedman test, * p < 0.05, ** p < 0.001).

**Supplementary Figure 5.**
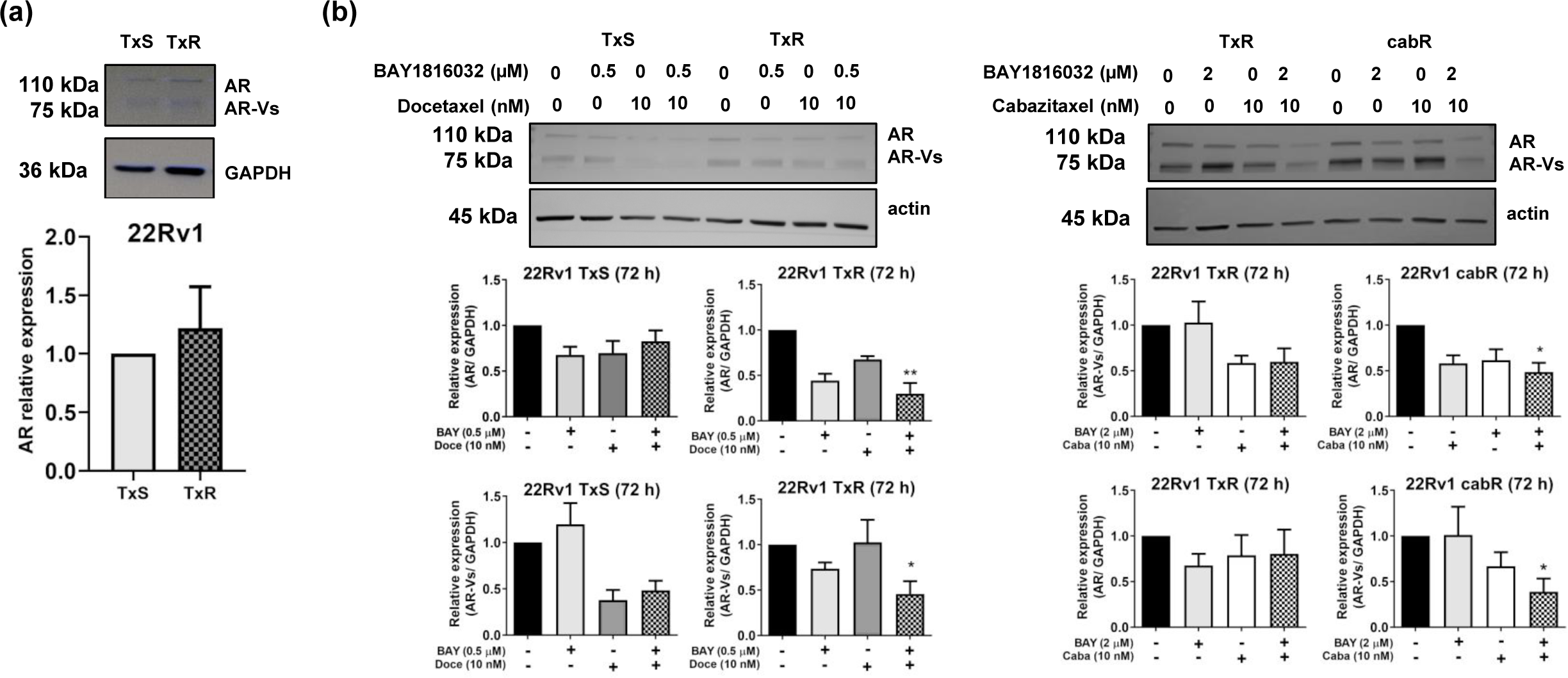
AR levels are not increased in taxane resistant CRPC models. **(a)** AR levels were measured in 22Rv1 TxS and TxR models by Western Blot. Graph represents the quantification of 4 independent experiments. **(b)** AR and AR-Vs expression levels were measured in 22Rv1 TxS, TxR and cabR cells after treatment with BAY 1816032 (0.5 or 2 µM), docetaxel (10 nM), cabazitaxel (10 nM) or their combination for 72 h. Graphs represent the quantification of four independent experiments. (Kruskal-Wallis test, * p < 0.05).

**Supplementary Table 1. IC50 calculation for docetaxel and cabazitaxel (a)** 22Rv1, PC3 and DU145 TxS, TxR and cabR cell lines were treated with different doses of docetaxel and cabazitaxel for 72 h. IC50 was calculated using the ATT Bioquest online IC50 calculator tool (https://www.aatbio.com/tools/ic50-calculator).

## References

1. Paschalis A, de Bono JS. Prostate Cancer 2020: “The Times They Are a’Changing”. Cancer Cell. 2020;38(1):25-7.

2. Desai K, McManus JM, Sharifi N. Hormonal Therapy for Prostate Cancer. Endocr Rev. 2021;42(3):354–73.

3. Siegel RL, Miller KD, Fuchs HE, Jemal A. Cancer statistics, 2022. CA Cancer J Clin. 2022;72(1):7-33.

4. Nader R, El Amm J, Aragon-Ching JB. Role of chemotherapy in prostate cancer. Asian J Androl. 2018;20(3):221–9.

5. Mita AC, Figlin R, Mita MM. Cabazitaxel: more than a new taxane for metastatic castrate-resistant prostate cancer? Clin Cancer Res. 2012;18(24):6574–9.

6. Lombard AP, Gao AC. Resistance Mechanisms to Taxanes and PARP Inhibitors in Advanced Prostate Cancer. Curr Opin Endocr Metab Res. 2020;10:16–22.

7. Maloney SM, Hoover CA, Morejon-Lasso LV, Prosperi JR. Mechanisms of Taxane Resistance. Cancers (Basel). 2020;12(11).

8. Shiota M, Dejima T, Yamamoto Y, Takeuchi A, Imada K, Kashiwagi E, et al. Collateral resistance to taxanes in enzalutamide-resistant prostate cancer through aberrant androgen receptor and its variants. Cancer Sci. 2018;109(10):3224–34.

9. Thadani-Mulero M, Portella L, Sun S, Sung M, Matov A, Vessella RL, et al. Androgen receptor splice variants determine taxane sensitivity in prostate cancer. Cancer Res. 2014;74(8):2270–82.

10. Shimizu Y, Tamada S, Kato M, Hirayama Y, Takeyama Y, Iguchi T, et al. Androgen Receptor Splice Variant 7 Drives the Growth of Castration Resistant Prostate Cancer without Being Involved in the Efficacy of Taxane Chemotherapy. J Clin Med. 2018;7(11).

11. Tagawa ST, Antonarakis ES, Gjyrezi A, Galletti G, Kim S, Worroll D, et al. Expression of AR-V7 and ARv(567es) in Circulating Tumor Cells Correlates with Outcomes to Taxane Therapy in Men with Metastatic Prostate Cancer Treated in TAXYNERGY. Clin Cancer Res. 2019;25(6):1880–8.

12. Gurioli G, Conteduca V, Brighi N, Scarpi E, Basso U, Fornarini G, et al. Circulating tumor cell gene expression and plasma AR gene copy number as biomarkers for castration-resistant prostate cancer patients treated with cabazitaxel. BMC Med. 2022;20(1):48.

13. Wang Z, Shen H, Ma N, Li Q, Mao Y, Wang C, et al. The Prognostic Value of Androgen Receptor Splice Variant 7 in Castration-Resistant Prostate Cancer Treated With Novel Hormonal Therapy or Chemotherapy: A Systematic Review and Meta-analysis. Front Oncol. 2020;10:572590.

14. Magani F, Bray ER, Martinez MJ, Zhao N, Copello VA, Heidman L, et al. Identification of an oncogenic network with prognostic and therapeutic value in prostate cancer. Mol Syst Biol. 2018;14(8):e8202.

15. Bolanos-Garcia VM, Blundell TL. BUB1 and BUBR1: multifaceted kinases of the cell cycle. Trends Biochem Sci. 2011;36(3):141–50.

16. Faesen AC, Thanasoula M, Maffini S, Breit C, Muller F, van Gerwen S, et al. Basis of catalytic assembly of the mitotic checkpoint complex. Nature. 2017;542(7642):498-502.

17. Zhu LJ, Pan Y, Chen XY, Hou PF. BUB1 promotes proliferation of liver cancer cells by activating SMAD2 phosphorylation. Oncol Lett. 2020;19(5):3506–12.

18. Jiang N, Liao Y, Wang M, Wang Y, Wang K, Guo J, et al. BUB1 drives the occurrence and development of bladder cancer by mediating the STAT3 signaling pathway. J Exp Clin Cancer Res. 2021;40(1):378.

19. Song J, Ni C, Dong X, Sheng C, Qu Y, Zhu L. bub1 as a potential oncogene and a prognostic biomarker for neuroblastoma. Front Oncol. 2022;12:988415.

20. Takeda M, Mizokami A, Mamiya K, Li YQ, Zhang J, Keller ET, et al. The establishment of two paclitaxel-resistant prostate cancer cell lines and the mechanisms of paclitaxel resistance with two cell lines. Prostate. 2007;67(9):955–67.

21. Machioka K, Izumi K, Kadono Y, Iwamoto H, Naito R, Makino T, et al. Establishment and characterization of two cabazitaxel-resistant prostate cancer cell lines. Oncotarget. 2018;9(22):16185–96.

22. Zhao N, Peacock SO, Lo CH, Heidman LM, Rice MA, Fahrenholtz CD, et al. Arginine vasopressin receptor 1a is a therapeutic target for castration-resistant prostate cancer. Sci Transl Med. 2019;11(498).

23. Grasso CS, Wu YM, Robinson DR, Cao X, Dhanasekaran SM, Khan AP, et al. The mutational landscape of lethal castration-resistant prostate cancer. Nature. 2012;487(7406):239-43.

24. Cai C, Wang H, He HH, Chen S, He L, Ma F, et al. ERG induces androgen receptor-mediated regulation of SOX9 in prostate cancer. J Clin Invest. 2013;123(3):1109–22.

25. Chen L, Song Y, Hou T, Li X, Cheng L, Li Y, et al. Circ_0004087 interaction with SND1 promotes docetaxel resistance in prostate cancer by boosting the mitosis error correction mechanism. J Exp Clin Cancer Res. 2022;41(1):194.

26. Dhital B, Santasusagna S, Kirthika P, Xu M, Li P, Carceles-Cordon M, et al. Harnessing transcriptionally driven chromosomal instability adaptation to target therapy-refractory lethal prostate cancer. Cell Rep Med. 2023;4(2):100937.

27. Shim M, Kim Y, Park Y, Ahn H. Taxane-based Chemotherapy Induced Androgen Receptor Splice Variant 7 in Patients with Castration-Resistant Prostate Cancer: A Tissue-based Analysis. Sci Rep. 2019;9(1):16794.

28. Jin W, Ye L. KIF4A knockdown suppresses ovarian cancer cell proliferation and induces apoptosis by downregulating BUB1 expression. Mol Med Rep. 2021;24(1).

29. Huang Z, Wang S, Wei H, Chen H, Shen R, Lin R, et al. Inhibition of BUB1 suppresses tumorigenesis of osteosarcoma via blocking of PI3K/Akt and ERK pathways. J Cell Mol Med. 2021;25(17):8442–53.

30. Moustakas A. The mitotic checkpoint protein kinase BUB1 is an engine in the TGF-beta signaling apparatus. Sci Signal. 2015;8(359):fs1.

31. Nyati S, Gregg BS, Xu J, Young G, Kimmel L, Nyati MK, et al. TGFBR2 mediated phosphorylation of BUB1 at Ser-318 is required for transforming growth factor-beta signaling. Neoplasia. 2020;22(4):163–78.

32. Nyati S, Schinske-Sebolt K, Pitchiaya S, Chekhovskiy K, Chator A, Chaudhry N, et al. The kinase activity of the Ser/Thr kinase BUB1 promotes TGF-beta signaling. Sci Signal. 2015;8(358):ra1.

33. Serrano-Del Valle A, Reina-Ortiz C, Benedi A, Anel A, Naval J, Marzo I. Future prospects for mitosis-targeted antitumor therapies. Biochem Pharmacol. 2021;190:114655.

34. Tischer J, Gergely F. Anti-mitotic therapies in cancer. J Cell Biol. 2019;218(1):10–1.

35. Cohen-Sharir Y, McFarland JM, Abdusamad M, Marquis C, Bernhard SV, Kazachkova M, et al. Aneuploidy renders cancer cells vulnerable to mitotic checkpoint inhibition. Nature. 2021;590(7846):486-91.

36. Stopsack KH, Whittaker CA, Gerke TA, Loda M, Kantoff PW, Mucci LA, et al. Aneuploidy drives lethal progression in prostate cancer. Proc Natl Acad Sci U S A. 2019;116(23):11390–5.

37. Quinton RJ, DiDomizio A, Vittoria MA, Kotynkova K, Ticas CJ, Patel S, et al. Whole-genome doubling confers unique genetic vulnerabilities on tumour cells. Nature. 2021;590(7846):492-7.

38. Al Nakouzi N, Cotteret S, Commo F, Gaudin C, Rajpar S, Dessen P, et al. Targeting CDC25C, PLK1 and CHEK1 to overcome Docetaxel resistance induced by loss of LZTS1 in prostate cancer. Oncotarget. 2014;5(3):667-78.

39. Shin SB, Woo SU, Yim H. Cotargeting Plk1 and androgen receptor enhances the therapeutic sensitivity of paclitaxel-resistant prostate cancer. Ther Adv Med Oncol. 2019;11:1758835919846375.

40. Giordano A, Liu Y, Armeson K, Park Y, Ridinger M, Erlander M, et al. Polo-like kinase 1 (Plk1) inhibition synergizes with taxanes in triple negative breast cancer. PLoS One. 2019;14(11):e0224420.

41. Hongo H, Kosaka T, Suzuki Y, Oya M. Discovery of a new candidate drug to overcome cabazitaxel-resistant gene signature in castration-resistant prostate cancer by in silico screening. Prostate Cancer Prostatic Dis. 2021.

42. Hongo H, Kosaka T, Suzuki Y, Mikami S, Fukada J, Oya M. Topoisomerase II alpha inhibition can overcome taxane-resistant prostate cancer through DNA repair pathways. Sci Rep. 2021;11(1):22284.

43. Baron AP, von Schubert C, Cubizolles F, Siemeister G, Hitchcock M, Mengel A, et al. Probing the catalytic functions of Bub1 kinase using the small molecule inhibitors BAY-320 and BAY-524. Elife. 2016;5.

44. Siemeister G, Mengel A, Fernandez-Montalvan AE, Bone W, Schroder J, Zitzmann-Kolbe S, et al. Inhibition of BUB1 Kinase by BAY 1816032 Sensitizes Tumor Cells toward Taxanes, ATR, and PARP Inhibitors In Vitro and In Vivo. Clin Cancer Res. 2019;25(4):1404–14.

45. Darshan MS, Loftus MS, Thadani-Mulero M, Levy BP, Escuin D, Zhou XK, et al. Taxane-induced blockade to nuclear accumulation of the androgen receptor predicts clinical responses in metastatic prostate cancer. Cancer Res. 2011;71(18):6019–29.

46. Yu B, Liu Y, Luo H, Fu J, Li Y, Shao C. Androgen receptor splicing variant 7 (ARV7) inhibits docetaxel sensitivity by inactivating the spindle assembly checkpoint. J Biol Chem. 2021;296:100276.

47. Zhang G, Liu X, Li J, Ledet E, Alvarez X, Qi Y, et al. Androgen receptor splice variants circumvent AR blockade by microtubule-targeting agents. Oncotarget. 2015;6(27):23358–71.

48. Antonarakis ES, Lu C, Luber B, Wang H, Chen Y, Nakazawa M, et al. Androgen Receptor Splice Variant 7 and Efficacy of Taxane Chemotherapy in Patients With Metastatic Castration-Resistant Prostate Cancer. JAMA Oncol. 2015;1(5):582–91.

49. Onstenk W, Sieuwerts AM, Kraan J, Van M, Nieuweboer AJ, Mathijssen RH, et al. Efficacy of Cabazitaxel in Castration-resistant Prostate Cancer Is Independent of the Presence of AR-V7 in Circulating Tumor Cells. Eur Urol. 2015;68(6):939–45.

